# Expression analysis and single-nucleotide polymorphisms of *MLPH* and *PMEL17* genes associated with melanin deposition in Xuefeng black-bone chickens

**DOI:** 10.1101/2021.10.08.463738

**Authors:** Dengyuying, Pengcanyang, Liuxu, Hechangqing, Guosongchang, Quxiangyong

## Abstract

Melanin deposition related genes such as *MLPH* and *PMEL17* play an important role in black-bone chicken. This study was aimed to identify and associate SNPs in the *MLPH* and *PMEL17* genes with melanin content of pectoral muscle (MCPM) in Xuefeng black-bone chicken. A total of 120 Xuefeng black-bone chickens at 120-day-old were randomly selected to measure blackness of pectoral muscle (BPM), according to the degree of BPM selected 22 high blackness (HB) and 22 low blackness (LB) chickens to determine the MCPM, and extract DNA and mRNA. The results indicated that the MCPM in the HB group was higher than in the LB group (*P* < 0.01), and the L value in the HB group was lower than in the LB group (*P* < 0.01). And we measured the mRNA expression levels of *MLPH* and *PMEL17* genes in pectoral muscle by quantitative real-time PCR. The results revealed that the mRNA expression levels of *MLPH* gene (*P* < 0.05) and *PMEL17* gene (*P* < 0.01) in the HB group was higher than in the LB group, and the mRNA relative expression level of *MLPH* and *PMEL17* genes with MCPM was positive correlation (*P* < 0.01). And the sequencing results found that a total of 17 SNPs were found in *MLPH* gene, the C-1411T was associated with MCPM (*P* < 0.05), there was no difference in MCPM among other locus (*P* > 0.05). And there were 10 SNPs in *PMEL17* gene, the G-1843C, C-2812T, and G-2794A were associated with MCPM (*P* < 0.05), there was no difference in the MCPM among other locus (*P* > 0.05). These SNPs could be molecular markers for breeding selection of blackness traits.

---

Blackness is an important economic trait of black-bone chicken. Consumers generally judge the quality of black-bone chicken based on blackness. The blackness darker, the chicken better. Therefore, in order to satisfy requirement of consumers, blackness trait has always been an important selection index in the breeding of black-bone chicken. However, the blackness depends on the melanin which is synthesized by melanocytes and deposited in nearby tissues. It has been revealed that melanin could scavenge free radicals (Bustamante *et al*. 1993), improve antioxidant capacity (Blarzino *et al*. 1999), provide photoprotection (Premi *et al*. 2015), and antibacterial (Nosanchuk & Casadevall 2003).

Melanin deposition is a complex process which is regulated by multiple genes, and the formation and transportation of melanosomes could affect melanin deposition, since melanosomes are the places where melanin is synthesized and stored (Lv *et al*. 2020). Melanophilin (*MLPH*) is a Rab27a and myosin Va binding protein that regulates melanosomes transport in melanocytes, which could accumulate melanosomes in melanocyte dendritic endings to affect melanosomes transfer from melanocytes to adjacent epidermal keratinocytes (Hume *et al*. 2007; Uyen *et al*. 2008). At present, the lightening of coat color caused by *MLPH* gene mutations has been discovered in animals, like rabbits (Lehner *et al*. 2013), quails (Bed’hom *et al*. 2012), and chickens (Vaez *et al*. 2008). It has also been reported that *PMEL17* gene is involved in melanosomes formation and transport in humans (Singh *et al*. 2008). Melanosomes development involves four steps. PMEL17, serves as the structural basis of fibrils in stage II, is mainly through proteolytic cleavage and forms amyloid fiber structure to maintain homeostasis within melanocytes, thereby in order to facilitate the synthesis and deposition of melanin in stage III and IV (Raposo *et al*. 2001; Slominski *et al*. 2004; Hellström *et al*. 2011). There have been a number of studies observed that *PMEL17* mutations could cause the change of coat or skin color in animals, such as mouse (Kwon *et al*. 1995), horse (Brunberg *et al*. 2006), cattle (Mészáros *et al*. 2015), and chicken (Kerje *et al*. 2004).

Xuefeng black-bone chicken, a valuable race native breed in the Hunan Province of China, is a dual-purpose chicken bred for its meat and eggs, which is loved by people due to its high nutritional value. At present, the blackness of Xuefeng black-bone chicken varies greatly among individuals. The conventional breeding methods are slow and cost, genetic marker could effectively speed up the breeding selection process of blackness traits. Therefore, the objective of this study was to screen SNP (single nucleotide polymorphism) related to melanin deposition and provide a theoretical basis for the breeding of black-bone chickens. We selected *MLPH* and *PMEL17* genes as candidate genes for this experiment, analyzed its mRNA expression level in different blackness chickens, and evaluated the relationship between the SNP in candidate gene and melanin deposition.

## 1. Materials and Methods

All the birds and the experimental protocols were approved by the Institutional Animal Care and Use Committee of Hunan Agricultural University, Hunan, China.

### 1.1 Chickens and sample collection

In this study, the Xuefeng black-bone chicken was obtained from Hunan Yunfeifeng Agricultural Commercial Company (Hunan, China). A total of 120 Xuefeng black-bone chickens at 120-day-old were randomly selected to measure blackness of pectoral muscle (BPM). According to the degree of BPM selected 22 high blackness (HB) and 22 low blackness (LB) chickens, and slaughtered (half male and female) them to get the pectoralis major tissue on the same side, meanwhile, put samples quickly in cryogenic vials with RNase-free equipment and then stored in liquid nitrogen. Within 24 h, the samples were transferred to −80°C storage for subsequent determination.

### 1.2 Determination of blackness and melanin content in pectoral muscle

The BPM of Xuefeng black-bone chicken (measured twice for each chicken) was measured with clorimeter (Hunter Associates Laboratory, Inc. Virginia, USA), which performs white board calibrated before the measurement. The measurement scale is set to L*a*b mode, and the L value represents the brightness of the sample, which range is 0 to 100. Determination of melanin content reference to Ozeki (Ozeki *et al*. 1995).

### 1.3 DNA extraction, RNA isolation and cDNA preparation

Genomic DNA was extracted by the Phenol/chloroform method (Sambrook et al. 1989), and total RNA was isolated from pectoral muscles sample using Trizol reagent (Invitrogen Life Technologies, Carlsbad, USA). The cDNA samples were prepared from 1 μL total RNA per reaction using a reverse transcription kit (Takara Biotechnology, Tokyo, Japan) and stored at - 20°C for qRT-PCR assays. The concentration and purity of genomic DNA and total RNA were determined by NanoDrop ND-2000 UV spectrophotometer (Thermo Fisher Scientific, Inc. Massachusetts, USA).

### 1.4 Polymerase chain reaction (PCR) primers and amplification

The primer pairs for the amplification of *MLPH, PEML17* genes, and *GAPDH* gene (served as an endogenous reference gene) were used as listed in Table 1, Full-length sequences of *MLPH* and *PMEL17* genes, 1000 bp upstream sequences and 1000 bp downstream sequences were obtained from GenBank. PCR primers were designed by Primer Premier 5.0 software, the primers were synthesized by Tsingke Biotechnology Co., Ltd (Beijing, China). The detailed information for PCR of *MLPH* and *PMEL17* genes has been presented in Table 2, respectively.

**Table 1.**
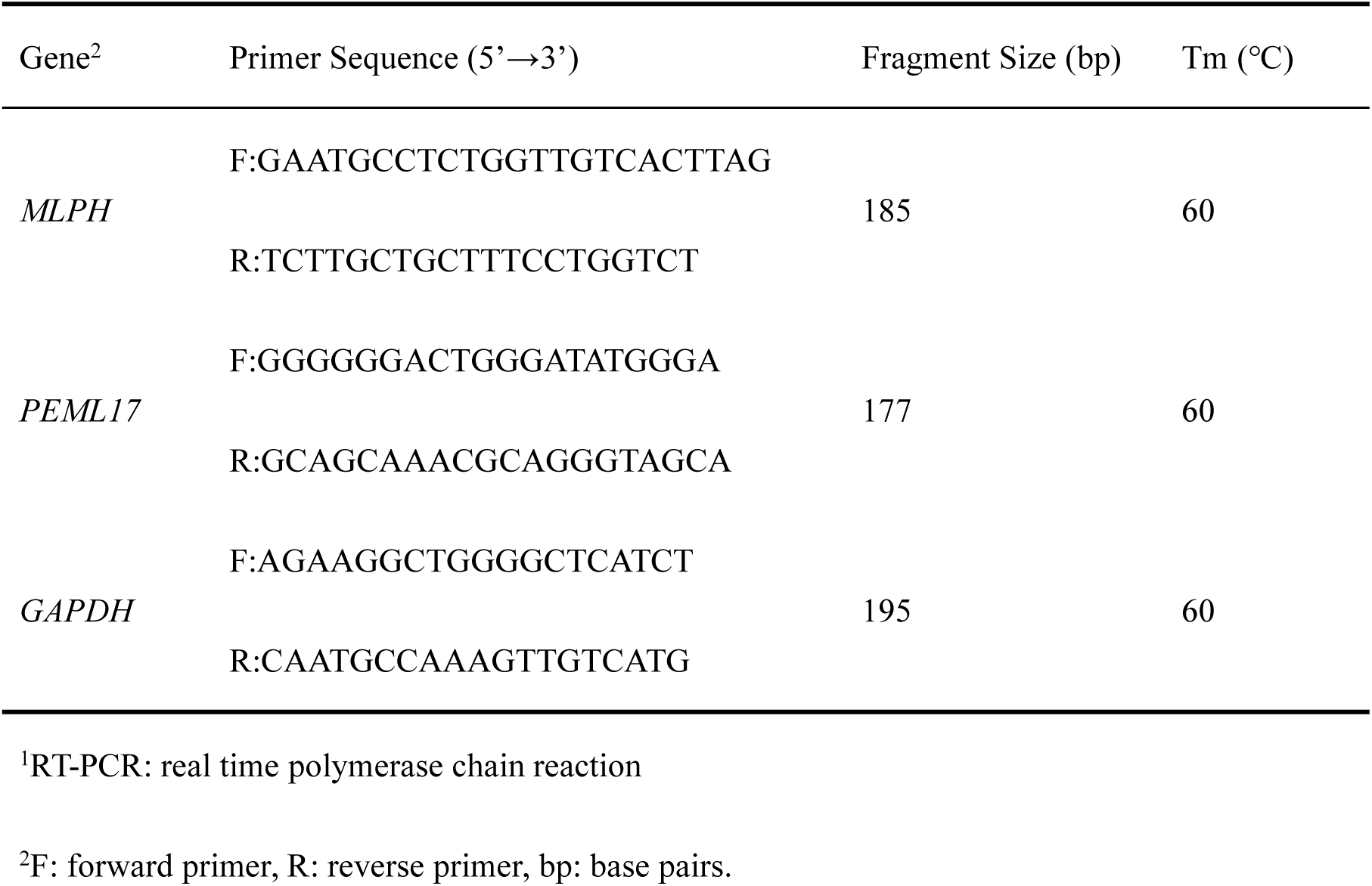
RT-PCR^1^ primer sequences

**Table 2.**
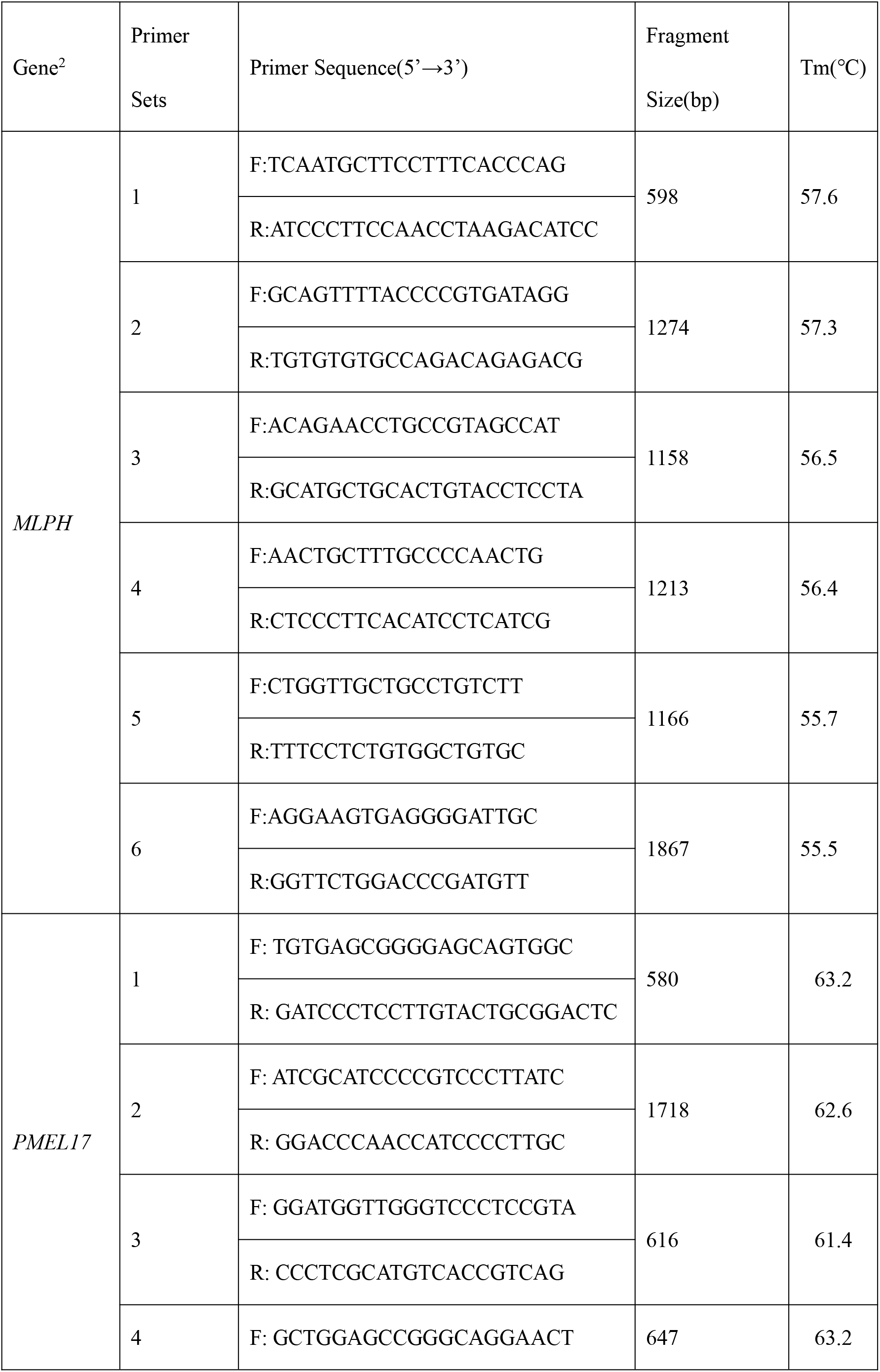
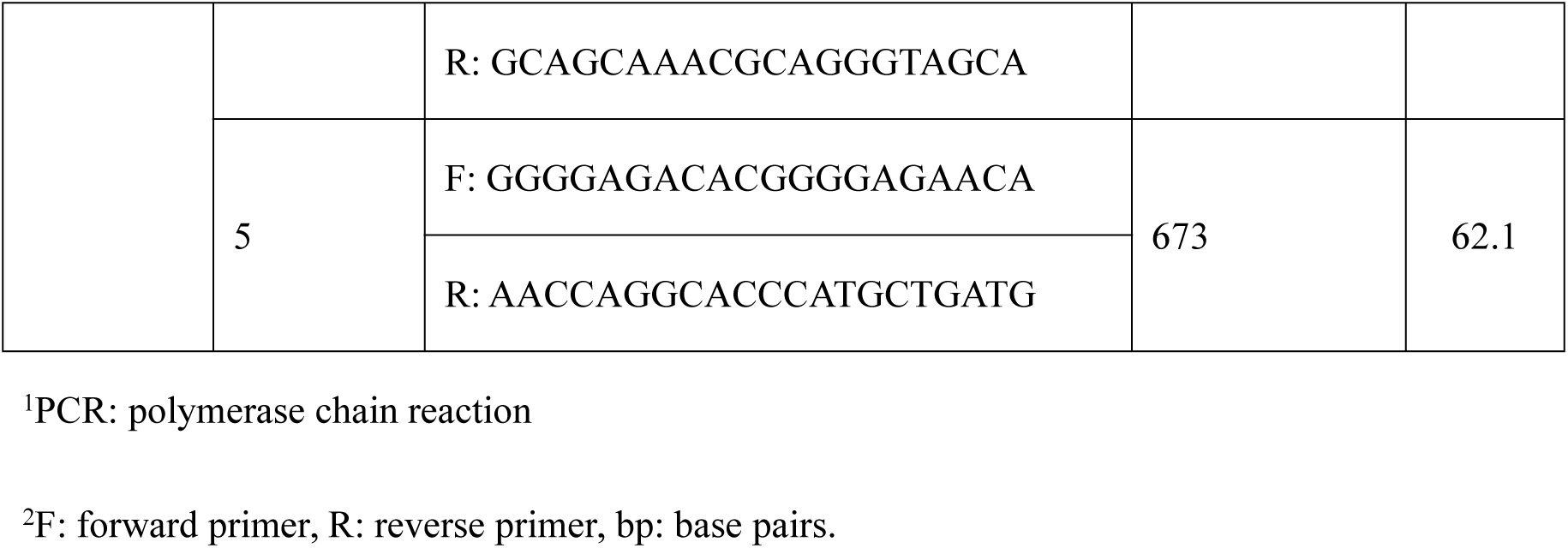
PCR^1^ primer sequences of *MLPH* and *PMEL17* genes

Using the SYBR Premix Ex TaqTM II kit (Takara Biotechnology, Tokyo, Japan) to perform the quantitative real-time PCR assay with CFX ConnectTM Real-Time PCR Detection System (Bio-Rad Laboratories, Hercules, California, USA). The RT-PCR amplification was carried out in 25 μL reaction volume, containing 2.0 μL cDNA, 1 μL of each primer, 12.5 μL of SYBR Premix Ex TaqTM (2×) and 8.5 μL double-distilled H_2_O. And the PCR amplification of *MLPH* and *PMEL17* genes was carried out in 20 μL reaction volume, containing 1.0 μL DNA, 0.8 μL of each primer, 12 μL of 2×Master mix and 5.4 μL double-distilled H_2_O.

### 1.4 Screening of SNPs in *MLPH* and *PMEL17* genes

The PCR products included all exon and some nearby intron sequences in *MLPH* and *PMEL17* genes, and sent it to Tsingke Biotechnology Co., Ltd (Beijing, China) for sequencing. The sequencing results were analyzed by Chromas and DNAStar software to screen out each SNPs site, and then sequenced a single sample and typed.

## 2. Statistical Analysis

The relative quantitative results were calculated using the 2^−ΔΔCt^ method after normalization against the reference gene *GAPDH*. The correlation analysis between the melanin content and the expression level of mRNA was performed by pearson correlation and significant differences among the melanin content of different genotype were analyzed using one-way ANOVA with duncans multiple comparison test, above all using SAS 9.2 version 21.0 statistical software (version 9.2, SAS Institute Inc., Cary, USA). All data are presented as mean ± standard error, considered statistically significant at *P*-Value less than 0.05 (*P* < 0.05).

Genotype frequencies and allelic frequencies were determined by direct counting, genotype frequency of SNPs was measured through chi-square test by Excel 2016, and checked it if consistent with the Hardy-Weinberg equilibrium. The least squares analysis was performed on the MCPM and carried out Tukey’s test, and significant level was set to be 0.05, the model used in the study was: Y=μ+G+e

Where: Y was the phenotype value of melanin, μ is the overall mean, G was the effect of genotype, and e was the effect of random residual errors.

## 3. Result

### 3.1 Correlation analysis of BPM and MCPM

Fig 1 showed a good linear relationship which is a melanin standard curve drawn by the absorbance of different concentrations of melanin standard solutions at 500 nm. According to Fig 2, the MCPM in HB group was higher than in LB group (*P* < 0.01), and the L value in HB group was lower than in LB group (*P* < 0.01).

**Fig 1:**
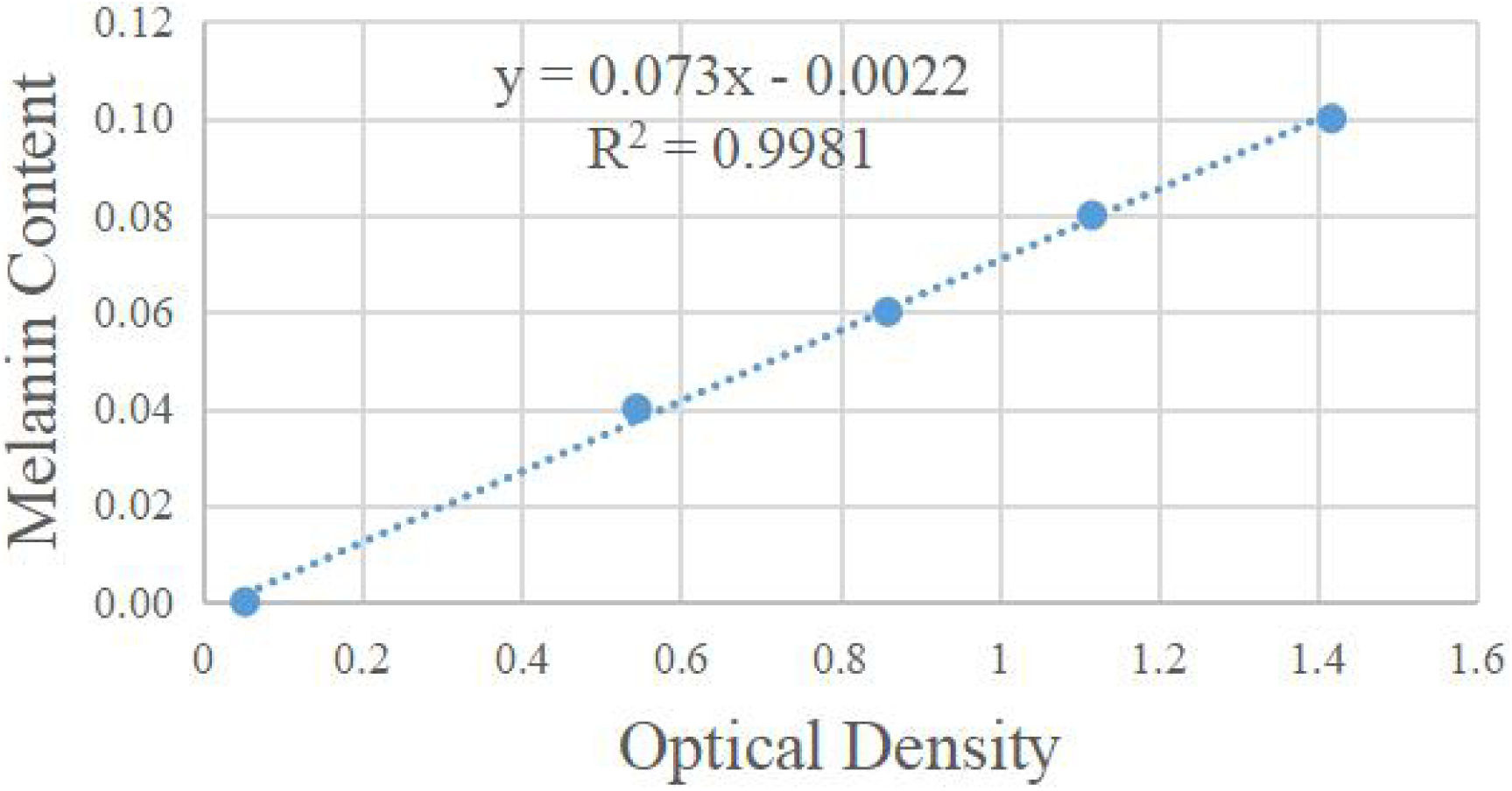
The standard curve of melanin

**Fig 2.**
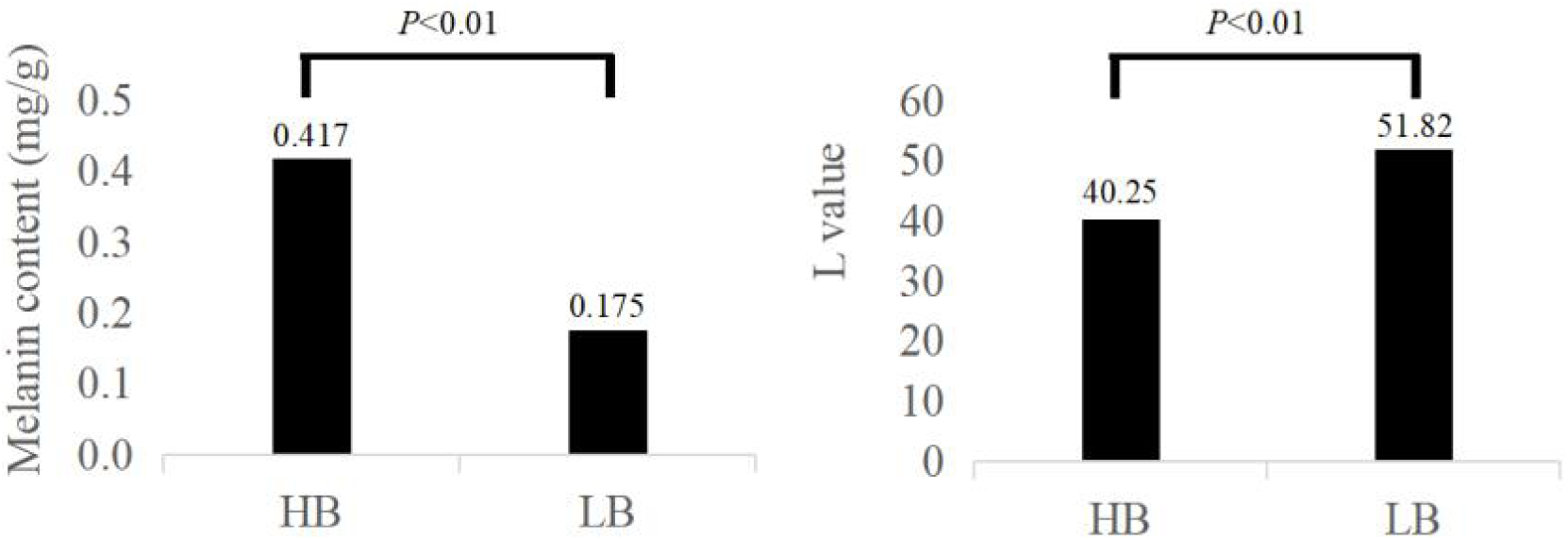
Melanin content, L value, W_1_ value, and W_2_ value in pectoral muscle^1^ ^1^HB: high blackness group; LB: low blackness group

### 3.2 Correlation analysis of the mRNA expression level of *MLPH, PMEL17* gene and MCPM

As showed in Fig 3-5, the mRNA relative expression level of *MLPH* (*P* < 0.05) and *PMEL17* (*P* < 0.01) in the HB group was higher than in the LB group. Through the correlation analysis found that the mRNA relative expression level of *MLPH* and *PMEL17* with MCPM was a positive correlation (*P* < 0.01). The results indicated that *MLPH* and *PMEL17* genes were closely related to MCPM of Xuefeng black-bone chicken.

**Fig 3.**
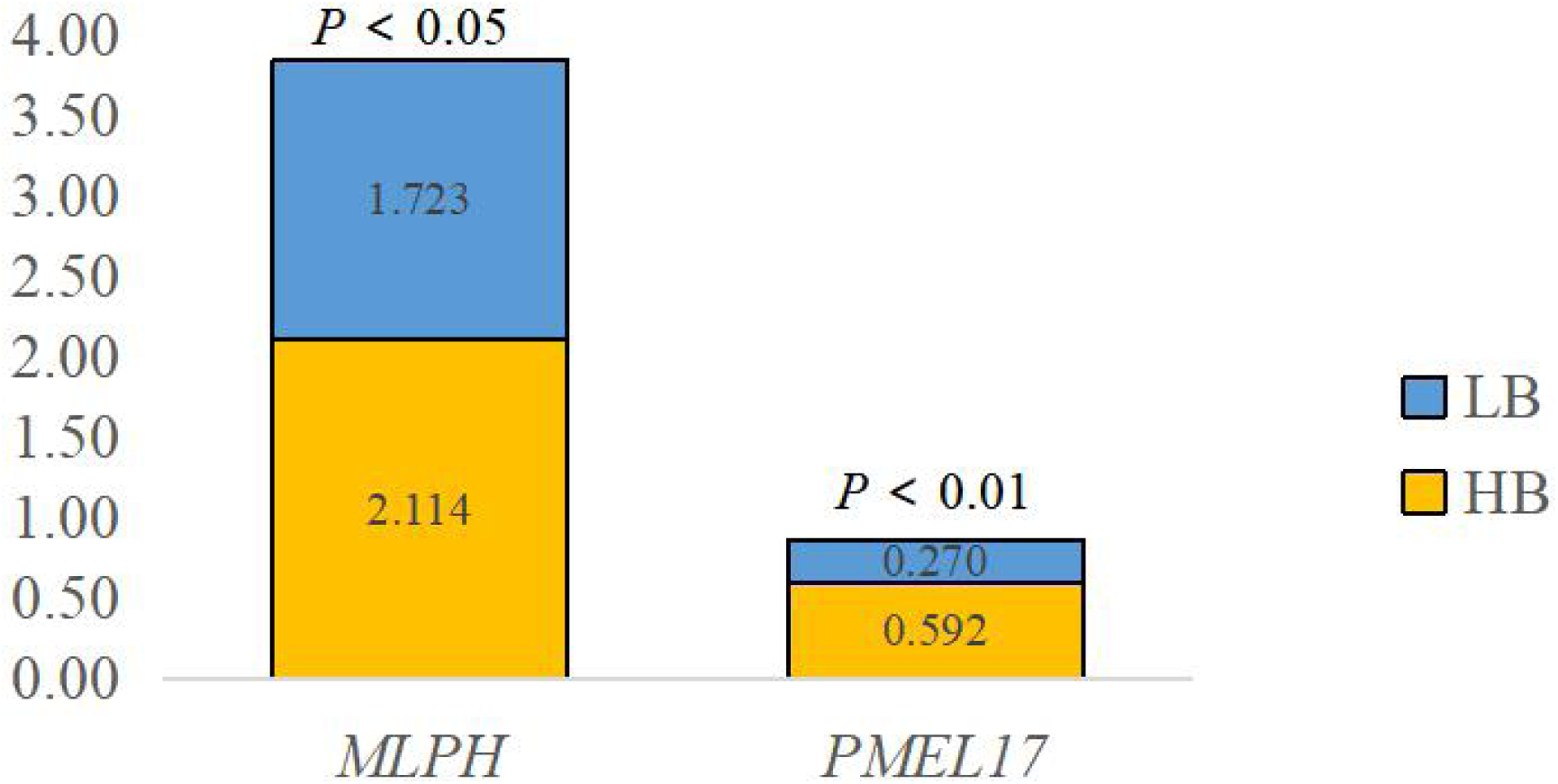
The mRNA relative expression of MLPH and PMEL 17 gene^1^ ^1^HB: high blackness group; LB: low blackness group

**Fig 4.**
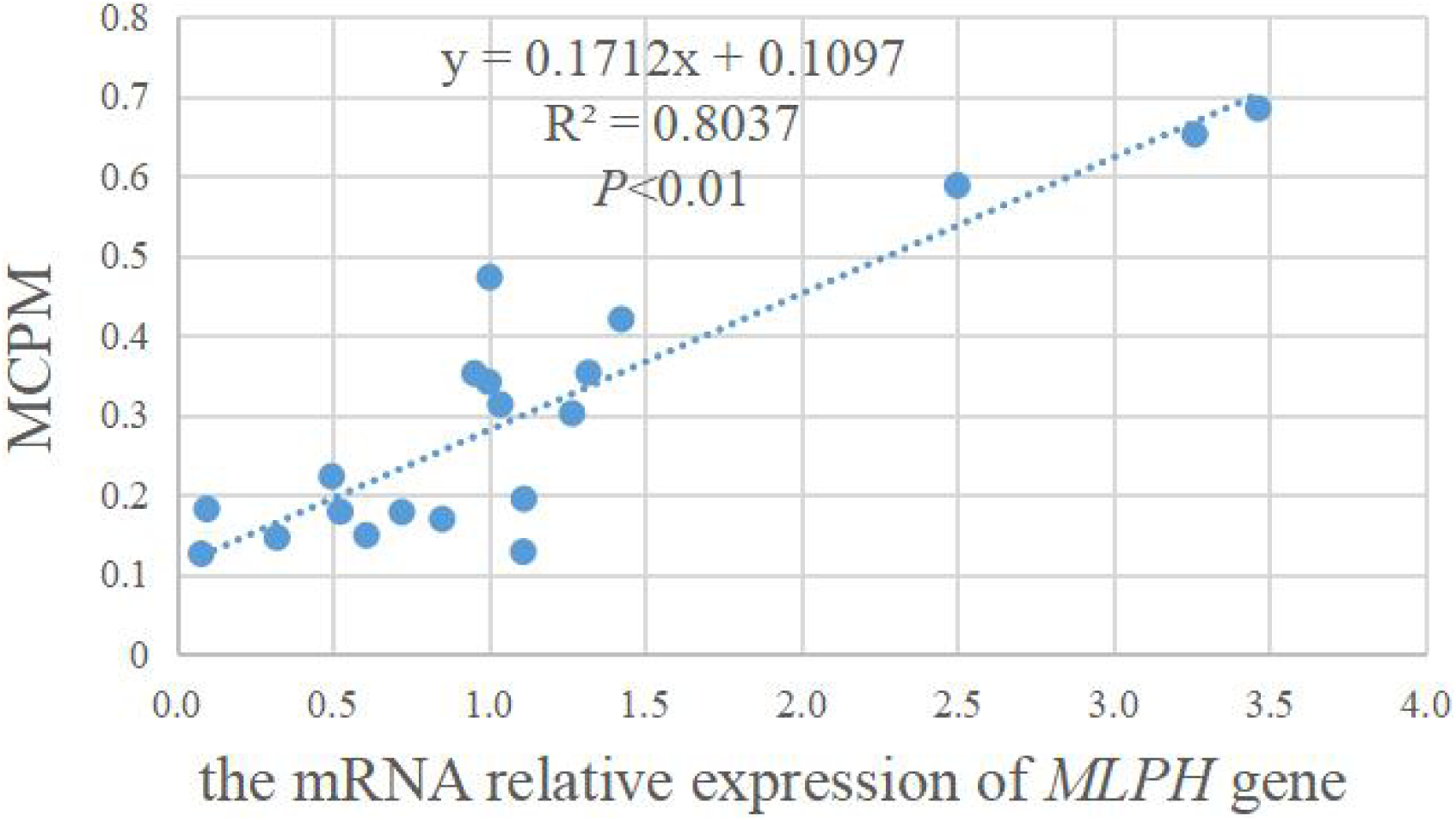
The linear relationship of relative expression of *MLPH* gene and MCPM^1^ ^1^ MCPM: melanin content of pectoral muscle

**Fig 5.**
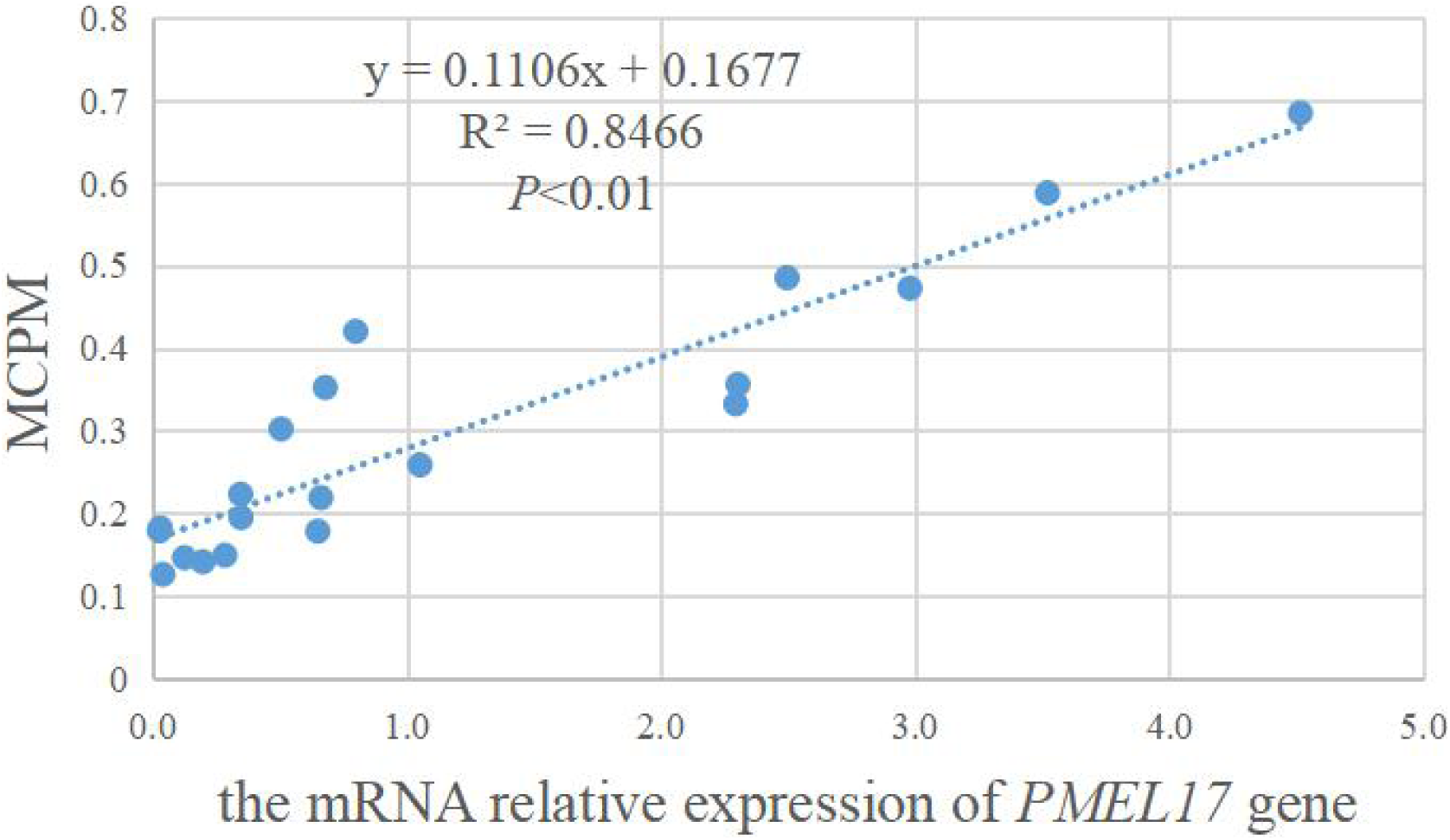
The linear relationship of the mRNA relative expression of *PMEL 17* gene and MCPM^1^ ^1^ MCPM: melanin content of pectoral muscle

### 3.3 Association of genetic variants of *MLPH, PMEL17* gene with MCPM

As presented in Fig 6-7, a total of 17 SNPs were found in the *MLPH* gene, 3 of which are in exon, and there were 10 SNPs in the *PMEL17* gene, 8 of which were in exon. All the SNPs in exon and their effects were summarized in Table 3. Allelic and genotypic distribution of the *MLPH* and *PMEL17* genes and MCPM in different genotypic were listed in Table 4-5, respectively. In the *MLPH* gene, the G-1414A were not in Hardy-Weinberg equilibrium (*P* < 0.05), and others were in Hardy-Weinberg equilibrium (*P* > 0.05). Meanwhile, the analysis revealed that the C-1411T was associated with MCPM (*P* < 0.05), and MCPM with genotype CC was higher than CT (*P* < 0.05). There was no difference in MCPM among other locus (*P* > 0.05). In the *PMEL17* gene, the G-2794A and G-3478A locus were not in Hardy-Weinberg equilibrium (*P* < 0.05), and others were in Hardy-Weinberg equilibrium (*P* > 0.05). And the analysis revealed that the G-1843C, C-2812T, and G-2794A locus were associated with MCPM (*P* < 0.05), there was no difference in MCPM among other locus (*P* > 0.05). In the G-1843C, MCPM of genotype GG was higher than GC and CC (*P* < 0.01), MCPM with genotype CC was higher than CT (*P* < 0.05) in the C-2812T, and MCPM with genotype AA was higher than GA and GG in the G-2794A (*P* < 0.05).

**Fig 6.**
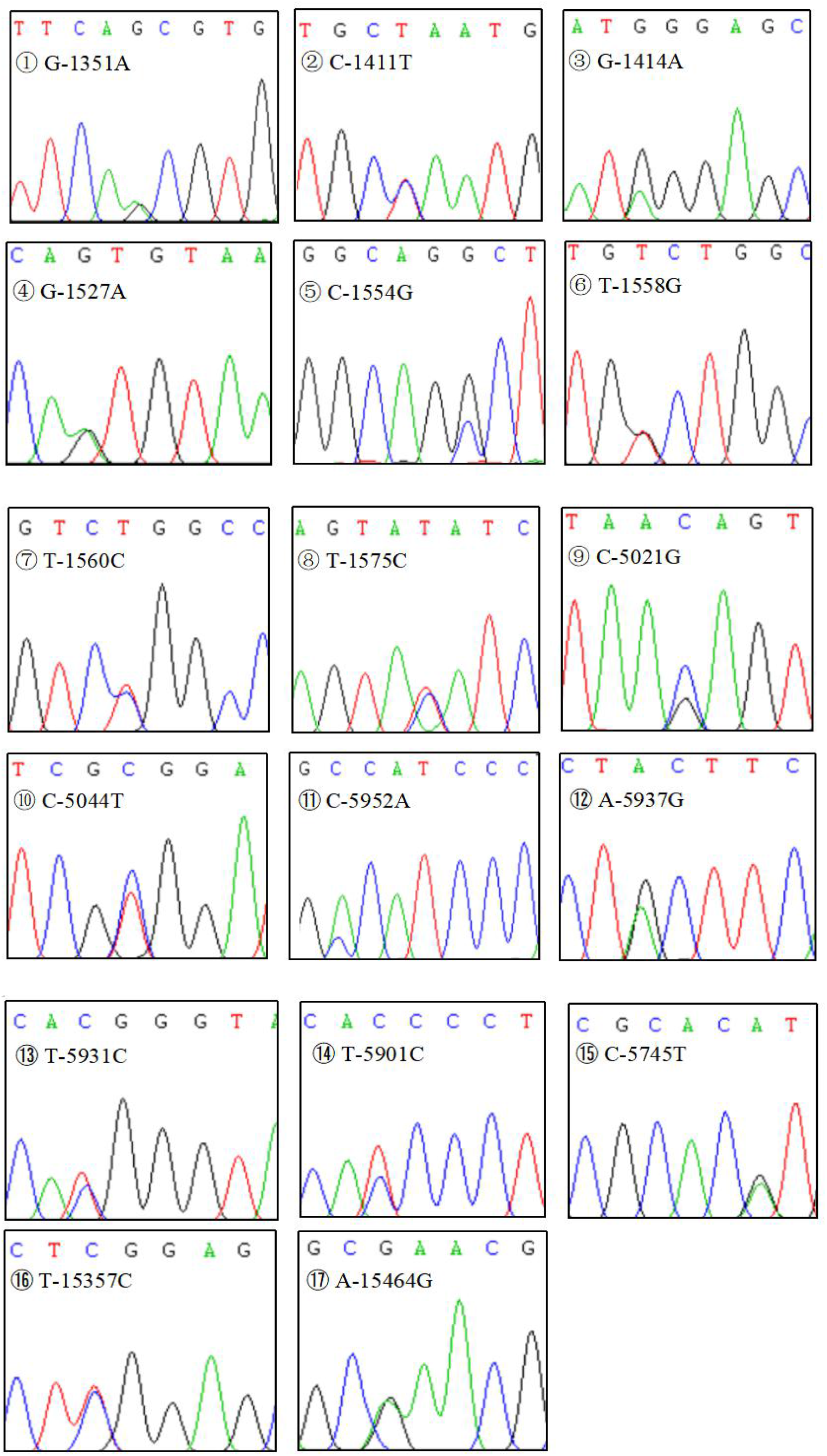
Chromatogram showing SNPs in *MLPH* gene

**Fig 7.**
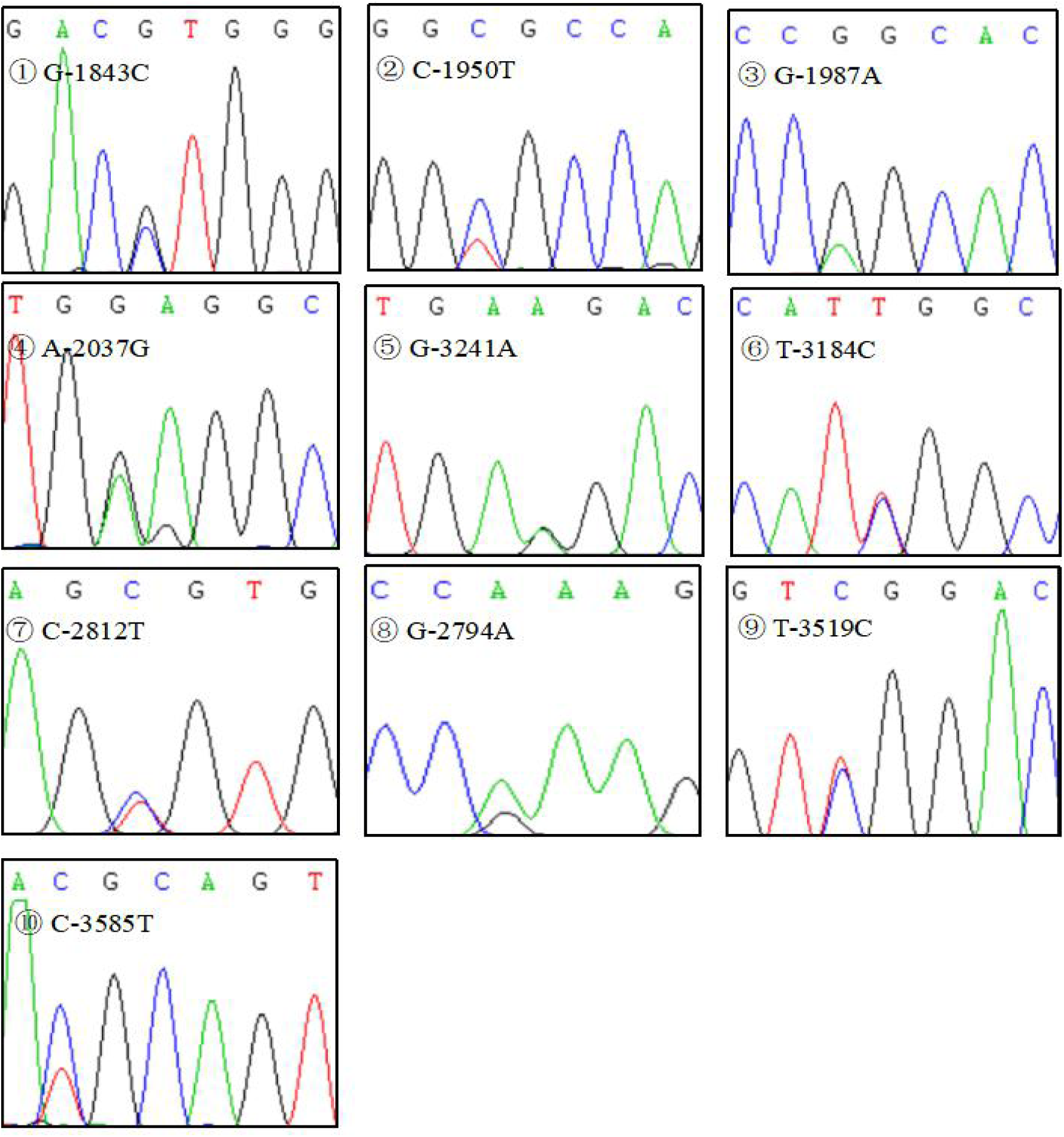
Chromatogram showing SNPs in *PMEL 17* gene

**Table 3.** Summary of SNPs identified in exon of *MLPH* and *PMEL17* genes

| Genes | SNPs | Type of variant | Change in amino acid |
| --- | --- | --- | --- |
| *MLPH* | G-1351A | synonymous variant | Glutamine |
|  | C-5745T | Missense variant | Arginine to Cystine |
|  | T-15357C |  | Leucine to Serine |
| *PMEL17* | T-3184C | synonymous variant | Proline |
|  | G-1987A |  |  |
|  | G-2794A | Missense variant | Serine to Leucine |
|  | C-2812T |  | Arginine to Histidine |
|  | G-3241A |  | Proline to Leucine |
|  | C-1950T |  | Alanine to Valine |
|  | T-3519C |  | Leucine to Serine |
|  | C-3585T |  | Threonine to Methionine |

**Table 4.** Allelic and genotypic distribution of the *MLPH* gene and MCPM in different genotypic^1^

| SNP | Genotype | Genotype frequency | Least-aquares | Allele | Allele frequency | Hardy-Weinberg equilibrium |  |
| --- | --- | --- | --- | --- | --- | --- | --- |
| | | | MCPM | | | $\chi^2$ value | *P* value |
| G-1351A | GG | 34 (0.773) | 0.312±0.027 | G | 0.886 | 0.72 | 0.395 |
|  | GA | 10 (0.227) | 0.229±0.028 | A | 0.114 |  |  |
|  | AA | 0 (0.000) |  |  |  |  |  |
| C-1411T | CC | 36 (0.818) | 0.314±0.026<sup>a</sup> | C | 0.909 | 0.44 | 0.507 |
|  | CT | 8 (0.182) | 0.196±0.029<sup>b</sup> | T | 0.091 |  |  |
|  | TT | 0 (0.000) |  |  |  |  |  |
| G-1414A | GG | 31 (0.705) | 0.281±0.029 | G | 0.807 | 5.20 | 0.023 |
|  | GA | 9 (0.205) | 0.358±0.042 | A | 0.193 |  |  |
|  | AA | 4 (0.091) | 0.238±0.055 |  |  |  |  |
| G-1527A | GG | 7 (0.159) | 0.308±0.064 | G | 0.409 | 0.05 | 0.821 |
|  | GA | 22 (0.500) | 0.300±0.032 | A | 0.591 |  |  |
|  | AA | 15 (0.341) | 0.275±0.041 |  |  |  |  |
| C-1554G | CC | 22 (0.500) | 0.305±0.036 | C | 0.716 | 0.17 | 0.683 |
|  | CG | 19 (0.432) | 0.284±0.033 | G | 0.284 |  |  |
|  | GG | 3 (0.068) | 0.258±0.072 |  |  |  |  |
| T-1558G | TT | 22 (0.500) | 0.305±0.036 | T | 0.716 | 0.47 | 0.495 |
|  | TG | 19 (0.432) | 0.284±0.033 | G | 0.284 |  |  |
|  | GG | 3 (0.068) | 0.258±0.072 |  |  |  |  |
| T-1560C | TT | 30 (0.682) | 0.274±0.025 | T | 0.845 | 1.37 | 0.241 |
|  | TC | 14 (0.318) | 0.334±0.047 | C | 0.155 |  |  |
|  | CC | 0 (0.000) |  |  |  |  |  |
| T-1575C | TT | 29 (0.659) | 0.275±0.029 | G | 0.807 | 0.12 | 0.729 |
|  | TC | 13 (0.295) | 0.327±0.043 | A | 0.193 |  |  |
|  | CC | 2 (0.045) | 0.328±0.043 |  |  |  |  |
| C-5021G | CC | 10 (0.227) | 0.244±0.039 | C | 0.489 | 0.09 | 0.760 |
|  | CG | 23 (0.523) | 0.333±0.035 | G | 0.511 |  |  |
|  | GG | 11 (0.250) | 0.254±0.038 |  |  |  |  |
| C-5044T | CC | 4 (0.091) | 0.392±0.036 | C | 0.261 | 0.60 | 0.437 |
|  | CT | 15 (0.341) | 0.345±0.047 | T | 0.739 |  |  |
|  | TT | 25 (0.568) | 0.246±0.025 |  |  |  |  |
| C-5952A | CC | 5 (0.114) | 0.353±0.061 | C | 0.364 | 0.28 | 0.594 |
|  | CA | 22 (0.500) | 0.313±0.033 | A | 0.636 |  |  |
|  | AA | 17 (0.386) | 0.249±0.036 |  |  |  |  |
| A-5937G | AA | 6 (0.114) | 0.353±0.061 | A | 0.364 | 0.28 | 0.594 |
|  | AG | 21 (0.500) | 0.313±0.033 | G | 0.636 |  |  |
|  | GG | 17 (0.386) | 0.249±0.036 |  |  |  |  |
| T-5931C | TT | 6 (0.114) | 0.353±0.061 | T | 0.364 | 0.28 | 0.594 |
|  | TC | 21 (0.500) | 0.313±0.033 | C | 0.636 |  |  |
|  | CC | 17 (0.386) | 0.249±0.036 |  |  |  |  |
| T-5901C | TT | 6 (0.114) | 0.353±0.061 | T | 0.364 | 0.28 | 0.594 |
|  | TC | 21 (0.500) | 0.313±0.033 | C | 0.636 |  |  |
|  | CC | 17 (0.386) | 0.249±0.036 |  |  |  |  |
| G-5745A | GG | 7 (0.159) | 0.337±0.061 | G | 0.455 | 1.62 | 0.204 |
|  | GA | 26 (0.591) | 0.295±0.031 | A | 0.545 |  |  |
|  | AA | 7 (0.250) | 0.259±0.042 |  |  |  |  |
| T-15357C | TT | 15 (0.341) | 0.255±0.022 | T | 0.625 | 1.98 | 0.159 |
|  | TC | 25 (0.568) | 0.303±0.035 | C | 0.375 |  |  |
|  | CC | 4 (0.091) | 0.370±0.103 |  |  |  |  |
| A-15464G | AA | 32 (0.727) | 0.308±0.029 | A | 0.830 | 3.37 | 0.066 |
|  | AG | 9 (0.205) | 0.289±0.036 | G | 0.170 |  |  |
|  | GG | 3 (0.068) | 0.136±0.043 |  |  |  |  |

<sup>1</sup> MCPM: melanin content of pectoral muscle, Means with different superscript differs
significantly

**Table 5.** Allelic and genotypic distribution of the *PMEL17* gene and MCPM in different genotypic^1^

| SNP | Genotype | Genotype frequency | Least-aquares | Allele | Allele frequency | Hardy-Weinberg equilibrium |  |
| --- | --- | --- | --- | --- | --- | --- | --- |
| | | | MCPM | | | $\chi^2$ value | *P* value |
| G-1843C | GG | 19 (0.432) | 0.376±0.036<sup>A</sup> | G | 0.614 | 2.39 | 0.122 |
|  | GC | 16 (0.364) | 0.239±0.034<sup>B</sup> | C | 0.386 |  |  |
|  | CC | 9 (0.205) | 0.211±0.028<sup>B</sup> |  |  |  |  |
| C-1950T | CC | 29 (0.659) | 0.266±0.025 | C | 0.807 | 0.12 | 0.729 |
|  | CT | 13 (0.295) | 0.367±0.048 | T | 0.193 |  |  |
|  | TT | 2 (0.045) | 0.192±0.066 |  |  |  |  |
| G-1987A | GG | 21 (0.477) | 0.343±0.033 | G | 0.693 | 0.01 | 0.920 |
|  | GA | 19 (0.432) | 0.247±0.035 | A | 0.307 |  |  |
|  | AA | 4 (0.091) | 0.251±0.043 |  |  |  |  |
| A-2037G | AA | 5 (0.114) | 0.425±0.061 | A | 0.250 | 3.27 | 0.070 |
|  | AG | 12 (0.273) | 0.285±0.037 | G | 0.750 |  |  |
|  | GG | 27 (0.614) | 0.272±0.030 |  |  |  |  |
| G-3241A | GG | 14 (13.642) | 0.301±0.048 | G | 0.557 | 0.05 | 0.827 |
|  | GA | 21 (21.716) | 0.322±0.034 | A | 0.443 |  |  |
|  | AA | 9 (8.642) | 0.212±0.016 |  |  |  |  |
| T-3184C | TT | 23 (0.523) | 0.276±0.029 | T | 0.716 | 0.11 | 0.739 |
|  | TC | 17 (0.386) | 0.311±0.041 | C | 0.284 |  |  |
|  | CC | 4 (0.091) | 0.315±0.090 |  |  |  |  |
| C-2812T | CC | 38 (0.864) | 0.315±0.025<sup>a</sup> | C | 0.932 | 0.24 | 0.627 |
|  | CT | 6 (0.136) | 0.152±0.011<sup>b</sup> | T | 0.068 |  |  |
|  | TT | 0 (0.000) |  |  |  |  |  |
| G-2794A | GG | 17 (0.386) | 0.216±0.023<sup>b</sup> | G | 0.523 | 9.05 | 0.030 |
|  | GA | 12 (0.273) | 0.346±0.048<sup>a</sup> | A | 0.477 |  |  |
|  | AA | 15 (0.341) | 0.338±0.043<sup>a</sup> |  |  |  |  |
| T-3519C | TT | 13 (0.295) | 0.264±0.041 | T | 0.511 | 0.81 | 0.367 |
|  | TC | 19 (0.432) | 0.340±0.035 | C | 0.489 |  |  |
|  | CC | 12 (0.273) | 0.249±0.041 |  |  |  |  |
| C-3585T | CC | 21 (0.477) | 0.282±0.033 | C | 0.648 | 2.82 | 0.093 |
|  | CT | 15 (0.341) | 0.300±0.037 | T | 0.352 |  |  |
|  | TT | 8 (0.182) | 0.309±0.067 |  |  |  |  |

<sup>1</sup> MCPM: melanin content of pectoral muscle, Means with different superscript differs
significantly

## 4. Discussion

The methods of determining melanin content include ultraviolet spectrophotometry and liquid chromatography, in this experiment, we chose the former. The result showed that the MCPM in the HB group was extremely significant higher than in the LB group and the L value in the HB group was extremely significantly lower than in the LB group. Hu et al and Du et al detected the melanin content at absorbance of A500, (Hu *et al*. 2019) reported that the melanin content of black Rex rabbit in skin was significantly higher than beaver, protein cyan, protein yellow, and white Rex rabbits. And (Du *et al*. 2017) found that the wild-type raccoon dog’s melanin content of guard hair and fluff was significantly higher than white-type. (Muroya *et al*. 2000) also demonstrated that melanin deposition was different in silky fowl different tissue, and melanin content of pectoralis in silky fowl was higher than white leghorn chicken. It reveals that melanin content could affect the visible blackness, which was similar to our results.

Melanin deposition is regulated by multiple genes which related more than 125 genes have been identified to date (Bennett & Lamoreux 2003). In this study, we chose two genes that affect the maturation and transport of melanosomes were selected as target genes for research. As a key structural protein of melanosome, *PMEL17* gene is mainly involved in the formation and transport of melanosomes (Singh *et al*. 2008; Sitaram & Marks 2012), while *MLPH* gene affects the transport of melanosomes to adjacent cells (Hume *et al*. 2007). The results indicated that the mRNA expression level of *MLPH* and *PMEL17* genes in the HB group was significantly higher than in the LB group. We speculate that *MLPH* and *PMEL17* genes may correlation with MCPM. Therefore, we have done further correlation analysis. The results indicated that the mRNA expression level of *MLPH* and *PMEL17* genes with MCPM was a significant positive correlation. There are a few reports on the mRNA expression of these two genes and their correlation with MCPM in black-bone chickens at present. and so as other animals. However, similar studies have shown that melanin-related genes are highly expressed in black phenotype, such as MITF (Lin *et al*. 2019), TYR (Anello *et al*. 2019), and MC1R (Li *et al*. 2019) genes. This suggested that *MLPH* and *PMEL17* genes are related to melanin deposition. However, the specific mechanism needs to be further studied.

SNPs, as molecular markers, have unique advantages in animals genetic breeding, and a single SNP could be directly used to research the association with related traits (Vandeputte *et al*. 2007; Ren *et al*. 2020). Missense mutation in exon could lead to amino acid sequences changes, which affect the structure and function of proteins, thereby causing phenotype differences between individuals (Cai *et al*. 2019; Bai *et al*. 2021). Meanwhile, the intron is involved in the splicing process of genes, which changes the coding of the amino acid through influencing the splicing efficiency or accuracy of gene translation, thereby altering the phenotype of individuals (Mayo 2008; Guo *et al*. 2016). This indicated that phenotype may be affected by multiple mutation locus at the same time. Research on Tianfu black rabbit and Sichuan white rabbit revealed that polymorphism distribution of three SNPs (c.693C < G, c.851A < G, c.911G < A) in the *MLPH* gene were significantly different (Jia *et al*. 2021). and Xu et al (Xu *et al*. 2016) tested the relationship between polymorphism of *MLPH* gene and the dilution phenotype to demonstrate that a mutation (C.1909A > G) in the *MLPH* gene causing the development of five-gray phenotype in the anyi tile-like gray chicken. There also have the same reported in dogs (Welle *et al*. 2009; Santos *et al*. 2017; Dudásová *et al*. 2021). In this paper, the results revealed that a total of 17 SNPs were found in *MLPH* gene, and 3 of which are in exon, and the C-1411T in intron had significant association with MCPM, which MCPM in genotype CC was significantly higher than CT.

Genotype TT was not detected, maybe it was eliminated in the artificial selection process and the C allele was more conducive to the melanin phenotype. And there were 10 SNPs in *PMEL17* gene, 8 of which were in exon. And the analysis revealed that the G-1843C, C-2812T, and G-2794A had significant association with MCPM. Previous studies have demonstrated that three different types of *PMEL17* gene mutations associated with plumage color in chickens (Kerje *et al*. 2004; Alhateemi *et al*. 2018). And it has also been reported in other animals that *PMEL17* gene mutation was related to phenotypic color changes (Brunberg *et al*. 2006; Hellström *et al*. 2011; Mészáros *et al*. 2015). In this study, one intron mutation (G-1843C) and two sense mutations (C-2812T and G-2794A) were found in *PMEL17* gene, and nonsense mutation in this gene can also cause phenotypic changes (Ishishita *et al*. 2018). This indicated that any mutation may change the phenotype. Meanwhile, three SNPs were in the Hardy-Weinberg equilibrium in the population, which indicated this SNPs have become stable in the long-term evolution and selection, and those SNPs could be molecular markers for the breeding of Xuefeng black-bone chicken.

## 5. Conclusion

In conclusion, the results indicate that *MLPH* and *PMEL17* genes are associated with melanin deposition in the pectoral muscle of Xuefeng black-boned chicken. And one SNP in *MLPH* gene and three SNPs in *PMEL17* gene were discovered, which could be as molecular markers for the melanin deposition in pectoral muscle. This study provides a theoretical basis for the breeding of Xuefeng black-boned chicken, and has enriched the information on breed protection and selection for local varieties.

## References

Alhateemi A.A.R., Sadeghi M., Shahrbabak M.M. & Shahrbabak H.M. (2018) Study of chicken PMEL17 gene polymorphism using DNA sequencing method in chicken and their effects on protein function % J Life Science Journal. 15.

Anello M., Fernández E., Daverio M.S., Vidal-Rioja L. & Di Rocco F. (2019) TYR Gene in Llamas: Polymorphisms and Expression Study in Different Color Phenotypes. Front Genet 10, 568.

Bai C., You Y., Liu X., Xia M., Wang W., Jia T., Pu T., Lu Y., Zhang C., Li X., Yin Y., Wang L., Zhou J. & Niu L. (2021) A novel missense mutation in the gene encoding major intrinsic protein (MIP) in a Giant panda with unilateral cataract formation. BMC Genomics 22, 100.

Bed’hom B., Vaez M., Coville J.L., Gourichon D., Chastel O., Follett S., Burke T. & Minvielle F. (2012) The lavender plumage colour in Japanese quail is associated with a complex mutation in the region of MLPH that is related to differences in growth, feed consumption and body temperature. BMC Genomics 13, 442.

Bennett D.C. & Lamoreux M.L. (2003) The color loci of mice--a genetic century. Pigment Cell Res 16, 333–44.

Blarzino C., Mosca L., Foppoli C., Coccia R., De Marco C. & Rosei M.A. (1999) Lipoxygenase/H2O2-catalyzed oxidation of dihdroxyindoles: synthesis of melanin pigments and study of their antioxidant properties. Free Radic Biol Med 26, 446–53.

Brunberg E., Andersson L., Cothran G., Sandberg K., Mikko S. & Lindgren G. (2006) A missense mutation in PMEL17 is associated with the Silver coat color in the horse. BMC Genet 7, 46.

Bustamante J., Bredeston L., Malanga G. & Mordoh J. (1993) Role of melanin as a scavenger of active oxygen species. Pigment Cell Res 6, 348–53.

Cai K., Wang J., Eissman J., Wang J., Nwosu G., Shen W., Liang H.C., Li X.J., Zhu H.X., Yi Y.H., Song J., Xu D., Delpire E., Liao W.P., Shi Y.W. & Kang J.Q. (2019) A missense mutation in SLC6A1 associated with Lennox-Gastaut syndrome impairs GABA transporter 1 protein trafficking and function. Exp Neurol 320, 112973.

Du Z., Huang K., Zhao J., Song X., Xing X., Wu Q., Zhang L. & Xu C. (2017) Comparative Transcriptome Analysis of Raccoon Dog Skin to Determine Melanin Content in Hair and Melanin Distribution in Skin. Sci Rep 7, 40903.

Dudásová S., Miluchová M. & Gábor M. (2021) Detection of MLPH gene polymorphism in population of the Belgian Shepherd variety Malinois. Journal of Central European Agriculture 22, 66–71.

Guo P., Zhao Z., Yan S., Li J., Xiao H., Yang D., Zhao Y., Jiang P. & Yang R. (2016) *PSAP* gene variants and haplotypes reveal significant effects on carcass and meat quality traits in Chinese Simmental-cross cattle. Archives Animal Breeding 59, 461–8.

Hellström A.R., Watt B., Fard S.S., Tenza D., Mannström P., Narfström K., Ekesten B., Ito S., Wakamatsu K., Larsson J., Ulfendahl M., Kullander K., Raposo G., Kerje S., Hallböök F., Marks M.S. & Andersson L. (2011) Inactivation of Pmel alters melanosome shape but has only a subtle effect on visible pigmentation. PLoS Genet 7, e1002285.

Hu S., Zhai P., Chen Y., Zhao B., Yang N., Wang M., Xiao Y., Bao G. & Wu X. (2019) Morphological Characterization and Gene Expression Patterns for Melanin Pigmentation in Rex Rabbit. Biochem Genet 57, 734–44.

Hume A.N., Ushakov D.S., Tarafder A.K., Ferenczi M.A. & Seabra M.C. (2007) Rab27a and MyoVa are the primary Mlph interactors regulating melanosome transport in melanocytes. J Cell Sci 120, 3111–22.

Ishishita S., Takahashi M., Yamaguchi K., Kinoshita K., Nakano M., Nunome M., Kitahara S., Tatsumoto S., Go Y., Shigenobu S. & Matsuda Y. (2018) Nonsense mutation in PMEL is associated with yellowish plumage colour phenotype in Japanese quail. Sci Rep 8, 16732.

Jia X., Ding P., Chen S., Zhao S., Wang J. & Lai S. (2021) Analysis of MC1R, MITF, TYR, TYRP1, and MLPH Genes Polymorphism in Four Rabbit Breeds with Different Coat Colors. Animals (Basel) 11.

Kerje S., Sharma P., Gunnarsson U., Kim H., Bagchi S., Fredriksson R., Schütz K., Jensen P., von Heijne G., Okimoto R. & Andersson L. (2004) The Dominant white, Dun and Smoky color variants in chicken are associated with insertion/deletion polymorphisms in the PMEL17 gene. Genetics 168, 1507–18.

Kwon B.S., Halaban R., Ponnazhagan S., Kim K., Chintamaneni C., Bennett D. & Pickard R.T. (1995) Mouse silver mutation is caused by a single base insertion in the putative cytoplasmic domain of Pmel 17. Nucleic Acids Res 23, 154–8.

Lehner S., Gähle M., Dierks C., Stelter R., Gerber J., Brehm R. & Distl O. (2013) Two-exon skipping within MLPH is associated with coat color dilution in rabbits. PLoS One 8, e84525.

Li J., Chen W., Wu S., Ma T., Jiang H.& Zhang Q. (2019) Differential expression of MC1R gene in Liaoning Cashmere goats with different coat colors. Anim Biotechnol 30, 273–8.

Lin R., Lin W., Zhou S., Chen Q., Pan J., Miao Y., Zhang M., Huang Z. & Xiao T. (2019) Integrated Analysis of mRNA Expression, CpG Island Methylation, and Polymorphisms in the MITF Gene in Ducks (Anas platyrhynchos). Biomed Res Int 2019, 8512467.

Lv J., Fu Y., Cao Y., Jiang S., Yang Y., Song G., Yun C. & Gao R. (2020) Isoliquiritigenin inhibits melanogenesis, melanocyte dendricity and melanosome transport by regulating ERK-mediated MITF degradation. Exp Dermatol 29, 149–57.

Mayo O. (2008) A century of Hardy-Weinberg equilibrium. Twin Res Hum Genet 11, 249–56.

Mészáros G., Petautschnig E., Schwarzenbacher H. & Sölkner J. (2015) Genomic regions influencing coat color saturation and facial markings in Fleckvieh cattle. Anim Genet 46, 65–8.

Muroya S., Tanabe R., Nakajima I. & Chikuni K. (2000) Molecular characteristics and site specific distribution of the pigment of the silky fowl. J Vet Med Sci 62, 391–5.

Nosanchuk J.D. & Casadevall A. (2003) The contribution of melanin to microbial pathogenesis. Cell Microbiol 5, 203–23.

Ozeki H., Ito S., Wakamatsu K. & Hirobe T. (1995) Chemical characterization of hair melanins in various coat-color mutants of mice. J Invest Dermatol 105, 361–6.

Premi S., Wallisch S., Mano C.M., Weiner A.B., Bacchiocchi A., Wakamatsu K., Bechara E.J., Halaban R., Douki T.& Brash D.E. (2015) Photochemistry. Chemiexcitation of melanin derivatives induces DNA photoproducts long after UV exposure. Science 347, 842–7.

Raposo G., Tenza D., Murphy D.M., Berson J.F. & Marks M.S. (2001) Distinct protein sorting and localization to premelanosomes, melanosomes, and lysosomes in pigmented melanocytic cells. J Cell Biol 152, 809–24.

Ren T., Zhang Z., Fu R., Yang Y., Li W., Liang J., Mo G., Luo W. & Zhang X. (2020) A 51 bp indel polymorphism within the PTH1R gene is significantly associated with chicken growth and carcass traits. Anim Genet 51, 568–78.

Sambrook J., Fritsch E.F. & Maniatis T. (1989) Molecular cloning: a laboratory manual.

Santos L.M., Messas N.B., Palumbo M.I.P. & et a. (2017) Identification of SNP c.-22G> A in the melanophilin gene from a dog with color dilution alopecia: case report. Arquivo Brasileiro de Medicina Veterinária e Zootecnia 69, 1503–7.

Singh S.K., Nizard C., Kurfurst R., Bonte F., Schnebert S. & Tobin D.J. (2008) The silver locus product (Silv/gp100/Pmel17) as a new tool for the analysis of melanosome transfer in human melanocyte-keratinocyte co-culture. Exp Dermatol 17, 418–26.

Sitaram A. & Marks M.S. (2012) Mechanisms of protein delivery to melanosomes in pigment cells. Physiology (Bethesda) 27, 85–99.

Slominski A., Tobin D.J., Shibahara S. & Wortsman J. (2004) Melanin pigmentation in mammalian skin and its hormonal regulation. Physiol Rev 84, 1155–228.

Uyen L.D.P., Nguyen D.H. & Kim E.-K. (2008) Mechanism of skin pigmentation. Biotechnology and Bioprocess Engineering 13, 383–95.

Vaez M., Follett S.A., Bed’hom B., Gourichon D., Tixier-Boichard M. & Burke T. (2008) A single point-mutation within the melanophilin gene causes the lavender plumage colour dilution phenotype in the chicken. BMC Genet 9, 7.

Vandeputte M., Dupont-Nivet M., Chavanne H. & Chatain B. (2007) A polygenic hypothesis for sex determination in the European sea bass Dicentrarchus labrax. Genetics 176, 1049–57.

Welle M., Philipp U., Rüfenacht S., Roosje P., Scharfenstein M., Schütz E., Brenig B., Linek M., Mecklenburg L., Grest P., Drögemüller M., Haase B., Leeb T. & Drögemüller C. (2009) MLPH Genotype--Melanin Phenotype Correlation in Dilute Dogs. v. 100.

Xu J.G., Xie M.G., Zou S.Y., Liu X.F., Li X.H., Xie J.F. & Zhang X.Q. (2016) Interactions of allele E of the MC1R gene with FM and mutations in the MLPH gene cause the five-gray phenotype in the Anyi tile-like gray chicken. Genet Mol Res 15.

